# Improved Metagenomic Analysis for All-Food-Sequencing with AFS-MetaCache2: Illumina vs. Nanopore

**DOI:** 10.64898/2025.12.18.694891

**Authors:** André Müller, Alexander Wichmann, Felix Kallenborn, S. Lukas Hellmann, Thomas Hankeln, Bertil Schmidt

**Affiliations:** Department of Computer Science, Johannes Gutenberg University, Mainz, 55099, Germany; Institute of Organismic and Molecular Evolution, Johannes Gutenberg University, Mainz, 55099, Germany; Nucleic Acids Core Facility, Johannes Gutenberg University, Mainz, 55099, Germany

**Keywords:** Next-generation sequencing, Third-generation sequencing, Long-read sequencing, Quantitative metagenomics, Species identification, Food authentication, Eukaryotic genomes, Big data

## Abstract

**Background:** All-Food-Sequencing (AFS) is a method for untargeted metagenomic analysis that allows for the detection and quantification of food ingredients. While this approach avoids some of the shortcomings of targeted PCR-based methods, its performance depends on sequencing technologies, taxonomic classification tools, and genomic reference databases.

**Results:** AFS-MetaCache2 implements an improved reference database construction mechanism compared to prior approaches. To demonstrate the effectiveness to AFS, we sequenced sausages composed of mammalian and avian species using both short-read (Illumina) and long-read (Oxford Nanopore Technologies) platforms. While both approaches reliably detect the main components, our comparison shows that long-read sequencing is superior in terms of both quantification accuracy and false positive rates. The evaluation of representative metagenomic tools (Kraken2+Bracken, KrakenUniq, AFS-MetaCache1) demonstrates that AFS-MetaCache2 yields the best accuracy and fastest database build times, while reducing peak main memory consumption. It thus allows for efficient scaling to large reference genome sets.

**Conclusion:** Our study suggests that deep sequencing of total genomic DNA from samples with heterogeneous taxon composition, using 3rd generation sequencing technology followed by metagenomic analysis with AFS-MetaCache2, is a valuable approach for bio-surveillance of food ingredients. Our software is available at https://github.com/muellan/metacache.

## Background

Ensuring safety and quality standards is of high importance to the food industry. However, errors and fraud in the production and wrong labeling of food have garnered political and media attention in recent years. This motivates the need for analytical methods that allow for accurate monitoring of food species composition, ideally spanning various kingdoms of life including animals, plants, bacteria, fungi, and viruses. Quantitative real-time polymerase chain reaction (qPCR) [1–3], droplet digital PCR (ddPCR) [4–6], and sequencing of species-specific DNA bar codes [7–10] are technologies for food control that are widely used in practice. However, they are limited by the number and phylogenetic diversity of target species within a single assay and thus are not suitable for broad-scale screening.

As an alternative technology, high-throughput sequencing of metagenomic DNA from food samples has the potential to simultaneously screen for a wide range of species. Although prior definition of possible target species is not needed, subsequent bioinformatic analysis based on comparisons to genomic databases is required to identify and quantify actual food components. Our original All-Food-Seq (AFS) approach [11] mapped Illumina sequencing reads to a (small) number of reference genomes using BWA, and then determined species composition and relative quantities based on a read counting procedure. This detected anticipated species in food products with quantification accuracy comparable to ddPCR and simultaneously identified unexpected species in an untargeted approach [12].

Application of this approach in practical settings faces two challenges;

1. storing large amounts of reference genomes in increasingly big databases, and
2. accurate bioinformatics solutions for mapping and quantification.

To address these issues, AFS-Metacache1 [13] proposed a *k*-mer-based exact matching approach to compare each read to a database of reference genomes. To gain efficiency, subsampling of *k*-mers based on minhashing is employed to reduce both memory consumption and database construction times. Nevertheless, when considering scaling to large reference genome collections, the associated databases may still consume hundreds of gigabytes or even terabytes of memory, which makes scaling challenging.

In addition, false positive read classifications might be caused by regions of DNA shared among multiple reference genomes due to sequence conservation. A number of prior studies [14–16] in the field of microbial metagenomics or marker gene sequencing based on Illumina short-read sequencing have shown that performance depends both on the taxonomic classifier and utilized reference databases. Recent work further indicted that the accuracy of microbial taxonomic classification and profiling methods can be further improved by using long-read sequencing [17–20]. However, similar studies of metagenomics with complex eukaryotic genomes have not been conducted so far.

In this paper we therefore address both the **accuracy** and the **scalability** challenge of *k*-mer based AFS. Our contributions are two-fold:

### Long read sequencing

We are the first to investigate the impact of longer sequencing reads to broad-scale identification and quantification of species composition in food. Our experimental results using a number of calibrator sausages of known species sequenced with both Illumina and Oxford Nanopore Technologies (ONT) platforms show that using long-read datasets yield lower false-positive rates and at the same time can provide higher quantification accuracy.

### Scalable database construction

We also address the drawback of high main memory consumption and long database construction times of contemporary classification tools by proposing a novel reference genome database construction scheme. It produces a taxonomy-aware partitioning that automatically assigns similar reference genomes to different partitions by minimizing bucket overflows. The corresponding implementation is provided in *AFS-MetaCache2*. Our performance evaluation shows that AFS-MetaCache2 yields lower false-positive rates and higher quantification accuracy compared to Kraken2+Bracken [21, 22], KrakenUniq [23, 24], and AFS-MetaCache1 [13], while reducing both database construction times and peak main memory consumption by taking advantage of modern multi-core CPUs. Therefore, our new approach enables scalability to growing genome collections which is needed for practical broad-scale food screening.

## Methods

### DNA sequencing of sausage calibration samples

Thirteen calibration sausage samples containing admixtures of cattle, chicken, pig, sheep, horse and turkey at defined amounts were produced by a professional butchery and provided by the Official Food Control Authority of the Canton Zürich, Switzerland (see Table 1) [1]. The samples were prepared for calibration of foodstuff detection methods and reflect three different recipes of sausage production:

**Table 1.** Meat composition of our utilized calibrator sausages.

| Name | Cattle | Sheep | Pig | Horse | Chicken | Turkey |
| --- | --- | --- | --- | --- | --- | --- |
| KGefLyo_A | 0.5% | 0.0% | 5.5% | 0.0% | 14.0% | 80.0% |
| KGefLyo_B | 2.0% | 0.0% | 4.0% | 0.0% | 36.0% | 58.0% |
| KGefLyo_C | 4.0% | 0.0% | 2.0% | 0.0% | 58.0% | 36.0% |
| KGefLyo_D | 5.5% | 0.0% | 0.5% | 0.0% | 80.0% | 14.0% |
| KLyo_A | 14.0% | 0.0% | 80.0% | 0.0% | 0.5% | 5.5% |
| KLyo_B | 36.0% | 0.0% | 58.0% | 0.0% | 2.0% | 4.0% |
| KLyo_C | 58.0% | 0.0% | 36.0% | 0.0% | 4.0% | 2.0% |
| KLyo_D | 80.0% | 0.0% | 14.0% | 0.0% | 5.5% | 0.5% |
| Kal_A | 1.0% | 9.0% | 35.0% | 55.0% | 0.0% | 0.0% |
| Kal_B | 9.0% | 1.0% | 55.0% | 35.0% | 0.0% | 0.0% |
| Kal_C | 25.0% | 25.0% | 25.0% | 25.0% | 0.0% | 0.0% |
| Kal_D | 35.0% | 55.0% | 9.0% | 1.0% | 0.0% | 0.0% |
| Kal_E | 55.0% | 35.0% | 1.0% | 9.0% | 0.0% | 0.0% |

*Kal A-E:* all meat sausage,

*KLyo A-D:* Lyoner style sausage (matrix of meat, rind and lard), and

*KGeflLyo A-D:* poultry Lyoner (matrix of meat and skin).

Total DNA was extracted out of 200 mg homogenized sausage samples using the Wizard Plus system (Promega, 117 Madison, USA) according to the manufacturer’s protocol.

### Oxford Nanopore sequencing

DNA concentration was quantified using a DS-11 FX+ spectrophotometer/fluorometer (DeNovix). DNA integrity was assessed using a Fragment Analyzer 5200 with the DNF-474-1000 kit (Agilent Technologies).

Libraries were prepared using the Native Barcoding Kit 24 V14 (SQK-NBD114.24; Oxford Nanopore Technologies) according to the manufacturer’s ligation sequencing gDNA native barcoding v14 protocol. Briefly, 400 ng per sample was subjected to DNA repair using Ultra™ II End Repair/dA-Tailing Module (New England Biolabs). Native barcodes were ligated using the NEB Blunt/TA Ligase Master Mix (New England Biolabs). Library concentration was measured using the Qubit dsDNA HS assay (Thermo Fisher Scientific). Barcoded samples were purified using 0.4 x AMPure XP Beads (Beckmann Coulter) and pooled equimolarly. Sequencing adapters were ligated to the pooled DNA using Quick T4 DNA Ligase (New England Biolabs) and the library was loaded onto a R10.4.1 PromethION flow cell (FLO-PRO114M). Sequencing was performed on a PromethION 2 Solo for 48 h.

For downstream analysis, super-accuracy (SUP) basecalling was performed using Guppy basecall server v7.1.4 with model *dna r*10.4.1 *e*8.2 400*bps sup*@*v*4.2.0 with demultiplexing and adapter trimming enabled and discarding low quality reads (PHRED *<*8). This resulted in 1.9 to 4.3 million reads per sample (median: 1.5 million) with median read length of 374 bp and maximum read length of 21 kbp.

### Illumina sequencing

Sequencing library preparation and sequencing were performed by a commercial provider (StarSEQ, Mainz, Germany). The Nextera DNA Library Preparation Kit (Illumina, San Diego, USA) was applied following the manufacturer’s instructions. Typically, 1 ng of total DNA was used. Sequencing was carried out on an Illumina MiSeq instrument using reagent kit v.2 in 150 bp paired-end mode. All datasets were quality checked, trimmed and filtered by using the FASTQC data evaluation software and the trimmomatic v0.33 trimming tool. This resulted in 0.195 to 1.659 million reads per sample (median: 0.8 million) with median read length of 126 bp and maximum read length of 136 bp.

Datasets for both Illumina and ONT sequencing have been submitted to the SRA database under the project name PRJEB34001.

### AFS-MetaCache2 Pipeline

Our pipeline can be separated into two distinct phases: *build* and *query*. AFS-MetaCache2 extends the pipeline of AFS-MetaCache1 (as presented in [13]) by introducing a novel method for scalable database construction in the build phase. Figure 1 provides an overview of both phases which are further outlined in the following.

**Fig. 1.**
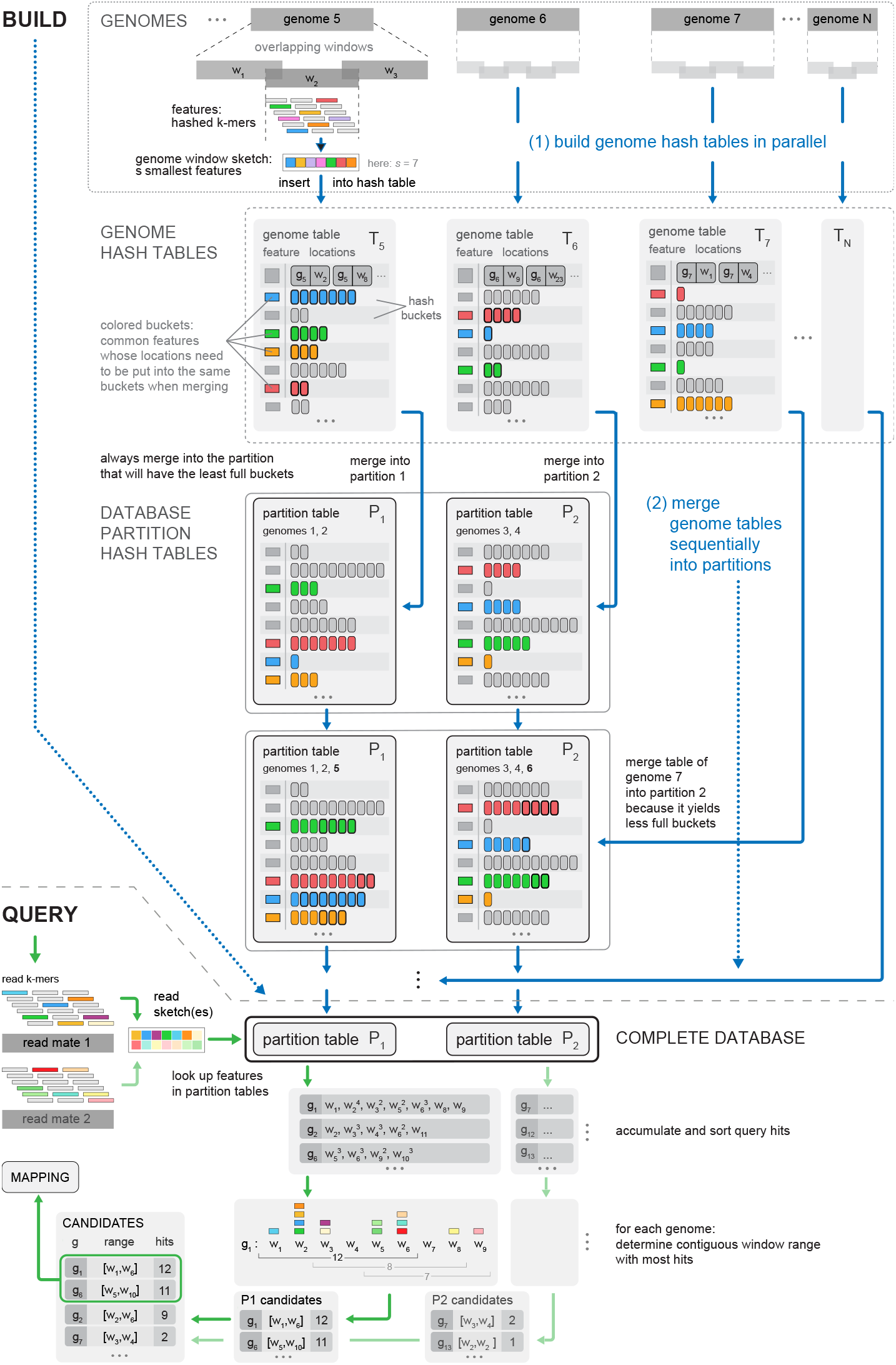
Top (build phase): Database building scheme. First, hash tables *T*_*i*_ are built in parallel for each reference genome *i*. Then, they are sequentially merged into the final database partitions *P*_*j*_ while minimizing overflowing hash buckets. Bottom (query phase): Querying pipeline. Features of each read (pair) are queried against each database partition and the resulting top candidates are merged to obtain the classification result.

### Build Phase

We consider a collection of *N* reference genomes as input. Each reference genome is divided into windows of size *l* which overlap by *k −* 1 base-pairs. For each window a sketch is calculated using minhashing [25]. A sketch consists of the *s* smallest *k*-mers (in strand-neutral canonical representation) contained in the window with respect to an applied hash function. Thus, the sketching procedure selects only a subset of *k*-mers to be inserted into the database used for similarity computation, typically reducing their amount by around one order-of-magnitude.

The database is stored as a hash table consisting of key-target-list pairs. A second hash function maps *k*-mers to slots in the hash table. If an identified slot is empty or occupied with the same *k*-mer, the corresponding *k*-mer is inserted as key and the corresponding location (genome ID, window ID) is appended to the target-list (see *Genome Hash Tables* in Figure 1).

Due to the ever increasing size of reference genome collections, the memory of a single workstation might not be sufficient to create a database for all considered reference sequences at once. Thus, we have developed a new approach to partition the reference genomes into separate databases (hash tables) which allows us to not only reduce memory consumption; but also limit the amount of overflowing buckets by storing similar genomes in different partitions. At the same time our approach reduces runtimes by employing multiple threads of common multi-core CPUs.

Our new database construction procedure for a total of *N* reference genomes *G*_1_, …, *G*_*N*_ works as follows. If *t* CPU threads are available in total, min(*t −* 1, *N*) individual hash tables *T*_*i*_ for each of the reference sequences *G*_*i*_ are constructed in parallel. They are then subsequently merged into larger hash tables using a separate, concurrently running thread. By default all genome-specific tables *T*_*i*_ are merged into one unified hash table which results in a database as produced by previous versions of AFS-MetaCache2. However, if the user either sets a maximum memory size per database partition or the number of individual partitions or both of these options, the genome-specific tables will be merged into separate hash tables each representing a database partition.

Since the distribution of *k*-mer occurrences in different genomes is usually highly skewed, a large fraction of *k*-mers occur only once while few occur thousands or even millions times. To limit memory consumption, the maximum number of locations stored per *k*-mer is limited to a predefined value (254 per default). In the case of highly similar genomes, the number of bucket collisions can become significant, since identical *k*-mers occur in several different genomes. This may lead to inaccurate assignments for reads stemming from repetitive genomic regions.

Thus, the merging of hash tables into multiple partitions *P*_*j*_ aims to preserve as much genomic information as possible by minimizing the number of overflowing hash table buckets. The procedure starts with a user-defined number of required partitions (by default 1) and successively inserts the contents of the previously generated per-genome hash tables *T*_*i*_ into these partitions. Each insertion round for any of the tables *T*_*i*_ starts by computing the number of resulting hash bucket overflows that would occur when inserting into any of the partitions *P*_*k*_. This can be achieved by querying each bucket key (*k*-mer) in *T*_*i*_ against *P*_*k*_ and in case of a match checking if the sum of the bucket sizes exceeds the maximum bucket size. Finally, the genome table is inserted into the partition *P*_*opt*_ that incurs the least amount of overflowing buckets and has the smallest size. If inserting the current genome table would exceed the maximum memory limit for partition *P*_*opt*_, the table *T*_*i*_ is not inserted but used to start a new partition instead.

Bucket truncation that could lead to a loss of information due to many *k*-mer collisions can thus be reduced significantly in our new build approach if two or more partitions are used. Moreover, it is not necessary to manually partition reference genomes in order to obtain size-limited partitions. Note that, as the partitioning works on a reference sequence level and *k*-mer distributions might differ between input sequences the resulting partitions might vary in size.

The parallel construction of individual sequence tables in AFS-MetaCache2 also significantly speeds up database construction by around one order-of-magnitude compared to prior methods such as AFS-MetaCache1, Kraken2, and KrakenUniq which are limited to a single thread operating a single hash table during the build phase. Note that merging incurs a small additional memory overhead since not yet merged individual tables *T*_*i*_ have to be kept alongside the partition tables which itself have to be kept large enough to accommodate new insertions, while not exceeding the maximum hash table load factor.

### Query Phase

To classify a read, its sequence is first split into windows of the same length as used in the database. From each window all canonical *k*-mers are generated and minhashing is applied to produce a sketch. All elements of the sketch are then queried against the hash table(s). The resulting location lists are merged and identical locations are accumulated. This yields a (sparse) histogram of hit counts per window in the reference genomes (window count statistic) which indicates the similarity of this region with the read. To account for single-end or paired-end Illumina reads spanning multiple windows, the window count statistic is scanned with a sliding window approach to find target regions with the highest aggregated hit counts in a contiguous window range. The top *m* counts (top hits) are then used to classify the read. In case of multiple partitions, each partition needs to be queried separately and the top-hit results need to be merged accordingly. If the difference of the highest and second highest count is above a threshold, the read is labeled as belonging to the taxon of the genome corresponding to the maximum count. Otherwise, all targets with counts close to the maximum are considered, the lowest common ancestor (LCA) of the corresponding taxa is calculated and used to label the read. Classifying ONT reads is identical to classifying (single-end) Illumina reads with the difference that ONT reads are typically longer and cover more windows.

Following the per-read classification, we perform *quantification* by estimating the abundances of organisms contained in a dataset at a specific taxonomical rank. For each taxon which occurs in the dataset we count the number of reads assigned to it. We then build a taxonomic tree containing all found taxa. Taxa on lower levels than the requested taxonomic rank are pruned and their read counts are added to their respective parents, while reads from taxa on higher levels are distributed among their children in proportion to the weights of the sub-trees rooted at each child. After the redistribution the estimated number of reads and abundance percentages are returned as outputs.

## Results

### Datasets

Sequencing read datasets have been obtained from thirteen calibrator sausage samples (Kal A-E, KLyo A-D, KGeflLyo A-D) with known meat composition (admixtures of chicken, turkey, pork, beef, horse, and sheep) as shown in Table 1. Table 2 provides a summary of the corresponding sequencing read datasets for both Illumina and ONT in terms of size and read length distribution. Due to the skewed read length distribution of ONT, we removed all reads shorter than 200-bp from the corresponding datasets.

**Table 2.** Description of utilized read datasets for each sequenced calibrator sausage for both Illumina and ONT.

| Dataset | Illumina |  |  | ONT |  |  |
| --- | --- | --- | --- | --- | --- | --- |
|  | Number of Seq. | Seq. Lengths Mean | Seq. Lengths Max | Number of Seq. | Seq. Lengths Mean | Seq. Lengths Max |
| KGefLyo_A | 675 648 | 123 | 136 | 3 730 696 | 483 | 13 550 |
| KGefLyo_B | 772 140 | 127 | 136 | 3 541 003 | 553 | 18 939 |
| KGefLyo_C | 703 550 | 128 | 136 | 3 606 972 | 509 | 14 817 |
| KGefLyo_D | 1 613 574 | 125 | 136 | 2 713 627 | 436 | 16 044 |
| KLyo_A | 801 976 | 128 | 136 | 1 878 926 | 580 | 21 061 |
| KLyo_B | 603 544 | 122 | 136 | 2 110 760 | 704 | 12 684 |
| KLyo_C | 1 014 268 | 125 | 136 | 2 617 000 | 674 | 21 048 |
| KLyo_D | 833 068 | 120 | 136 | 2 873 895 | 516 | 14 454 |
| Kal_A | 1 659 486 | 126 | 136 | 2 844 028 | 387 | 6 138 |
| Kal_B | 195 360 | 132 | 136 | 4 341 385 | 412 | 9 142 |
| Kal_C | 808 574 | 131 | 136 | 3 306 965 | 372 | 8 094 |
| Kal_D | 805 542 | 126 | 136 | 4 154 567 | 473 | 11 428 |
| Kal_E | 577 428 | 130 | 136 | 3 750 967 | 411 | 9 507 |

Food-related genomes (selection of main ingredients) used for database construction are listed in Table 3. We created databases of various sizes using the following reference genomes sets in order to compare quantification accuracy, runtime, and memory consumption of various tools:

**Table 3.** List of considered eukaryotic reference genomes.

| Item | Name | Accession ID | Size on disk |
| --- | --- | --- | --- |
| 01 | *Sus scrofa* (Pig) | GCF_000003025.6 | 2.4 GB |
| 02 | *Equus caballus* (Horse) | GCF_002863925.1 | 2.4 GB |
| 03 | *Meleagris gallopavo* (Turkey) | GCF_000146605.3 | 1.1 GB |
| 04 | *Mus musculus* (House Mouse) | GCF_000001635.26 | 2.7 GB |
| 05 | *Gallus gallus* (Chicken) | GCF_000002315.5 | 1.1 GB |
| 06 | *Ovis aries* (Sheep) | GCF_002742125.1 | 2.8 GB |
| 07 | *Rattus norvegicus* (Norway rat) | GCF_000001895.5 | 2.8 GB |
| 08 | *Bos taurus* (Cattle) | GCF_002263795.1 | 2.6 GB |
| 09 | *Bubalus bubalis* (Water buffalo) | GCF_003121395.1 | 2.6 GB |
| 10 | *Cervus elaphus hippelaphus* (Red deer) | GCA_002197005.1 | 3.3 GB |
| 11 | *Capreolus capreolus* (Western roe deer) | GCA_000751575.1 | 3.0 GB |
| 12 | *Struthio camelus australis* (African ostrich) | GCF_000698965.1 | 1.2 GB |
| 13 | *Anas platyrhynchos* (Mallard) | GCF_003850225.1 | 1.1 GB |
| 14 | *Capra hircus* (Goat) | GCF_001704415.1 | 2.8 GB |
| 15 | *Oryctolagus cuniculus* (Rabbit) | GCF_000003625.3 | 2.6 GB |
| 16 | *Cavia aperea* (Guinea pig) | GCA_000688575.1 | 2.6 GB |
| 17 | *Camelus ferus* (Wild batrian camel) | GCF_009834535.1 | 2.0 GB |
| 18 | *Canis lupus familiaris* (Dog) | GCF_000002285.3 | 2.3 GB |
| 19 | *Felis catus* (Domestic Cat) | GCF_000181335.3 | 2.4 GB |
| 20 | *Homo sapiens* (Human) | GCF_000001405.39 | 3.1 GB |
| 21 | *Equus asinus* (Donkey) | GCF_001305755.1 | 2.3 GB |
| 22 | *Rangifer tarandus* (Reindeer) | GCA_004026565.1 | 2.9 GB |
| 23 | *Phasianus colchicus* (Ring necked pheasant) | GCF_004143745.1 | 989 MB |
| 24 | *Glycine max* (Soybean) | GCF_000004515.5 | 946 MB |
| 25 | *Zea mays* (Maize) | GCF_902167145.1 | 2.1 GB |
| 26 | *Triticum aestivum* (Bread wheat) | GCA_002220415.3 | 15 GB |
| 27 | *Secale cereale* (Rye) | GCA_900079665.1 | 1.8 GB |
| 28 | *Hordeum vulgare* (Barley) | GCA_903813605.1 | 4.1 GB |
| 29 | *Oryza sativa japonica* (Rice) | GCF_000005425.2 | 370 MB |
| 30 | *Arachis hypogaea* (Peanut) | GCF_003086295.2 | 2.5 GB |
| 31 | *Saccharomyces cerevisiae* (Bakers yeast) | GCF_000146045.2 | 12 MB |
| 32 | *Notamacropus eugenii* (Tamar wallaby) | GCA_000004035.1 | 3.0 GB |
| 33 | *Brassica nigra* (Black mustard) | GCA_001682895.1 | 389 MB |
| 34 | *Brassica juncea* (Brown mustard) | GCA_001687265.1 | 924 MB |
| 35 | *Apium graveolens* (Celery) | GCA_009905375.1 | 3.2 GB |
| 36 | *Corylus avellana* (Hazelnut) | GCA_901000735.1 | 515 MB |
| 37 | *Sesamum indicum* (Sesame) | GCF_000512975.1 | 268 MB |
| 38 | *Juglans regia* (Walnut) | GCF_001411555.2 | 557 MB |
| 39 | *Pistacia vera* (Pistachio) | GCF_008641045.1 | 649 MB |
| 40 | *Prunus dulcis* (Almond) | GCF_902201215.1 | 220 MB |
| 41 | *Lupinus albus* (White lupine) | GCA_010261695.1 | 540 MB |
| 42 | *Lupinus angustifolius* (Blue lupine) | GCF_001865875.1 | 591 MB |
| Total |  |  | 89 GB |

*AFS8:* Genomes from 1 to 8 in Table 3.

*AFS20:* Genomes from 1 to 20 in Table 3.

*AFS31:* Genomes from 1 to 31 in Table 3.

*AFS42:* Genomes from 1 to 42 in Table 3.

To test the performance of AFS-MetaCache2 with partitioned databases, we built several version of AFS42 using one (AFS42), two (AFS42-P2), and four partitions (AFS42-P4).

### Quantification accuracy

Read classification and abundance estimation was performed for all sequencing read datasets listed in Table 2 using AFS-MetaCache2, Kraken2 v2.1.5 in combination with Bracken v2.5.3, and KrakenUniq v1.0.4 – all using default settings. AFS-MetaCache2 and Kraken2+Bracken were run with all eukaryotic databases of various sizes. KrakenUniq was only run with the AFS8 database, because it was not able to successfully build larger databases on our test system with 512 GB main memory.

In order to assess the overall performance of a mapping tool in combination with a given reference genome set and sequencing technology we use two metrics:

#### Absolute deviations

We sum up the absolute deviations from the known meat composition percentages for each dataset and compute the averages and standard deviations of these deviation sums over all datasets.

#### False positives

Read assignments to taxa other than the known main meat components are presumed to be false positive classifications^1^.

The results for absolute deviations are presented in Figure 2 which compares the average sums and mean deviations per dataset for AFS-MetaCache2, Kraken2+Bracken, and KrakenUniq. AFS-MetaCache2 yields consistently lower ground truth deviations than Kraken2+Bracken for all genome databases and both sequencing technologies. While KrakenUniq’s abundance results for AFS8 is comparable to AFS-MetaCache2 for the same database, it is not able to reliably scale to bigger genome collections. Deviation sums obtained with AFS-MetaCache2 range from (11.7± 3.1)% to (13.8 ± 2.6)% for Illumina reads and from (11.1± 3.4)% to (12.1 ± 2.4)% for ONT reads, while the corresponding values for Kraken2+Bracken range from (15.7 ±4.9)% to (16.3± 4.8)% for Illumina reads and from (14.8± 3.4)% to (16.3 ± 4.6)% for ONT reads. The deviation sums for KrakenUniq with the AFS8 database are (12.7± 3.1)% for Illumina and (12.6 ± 2.8)% for ONT respectively. Each tested tool thus achieves better average abundance accuracy for ONT reads than for Illumina reads.

**Fig. 2.**
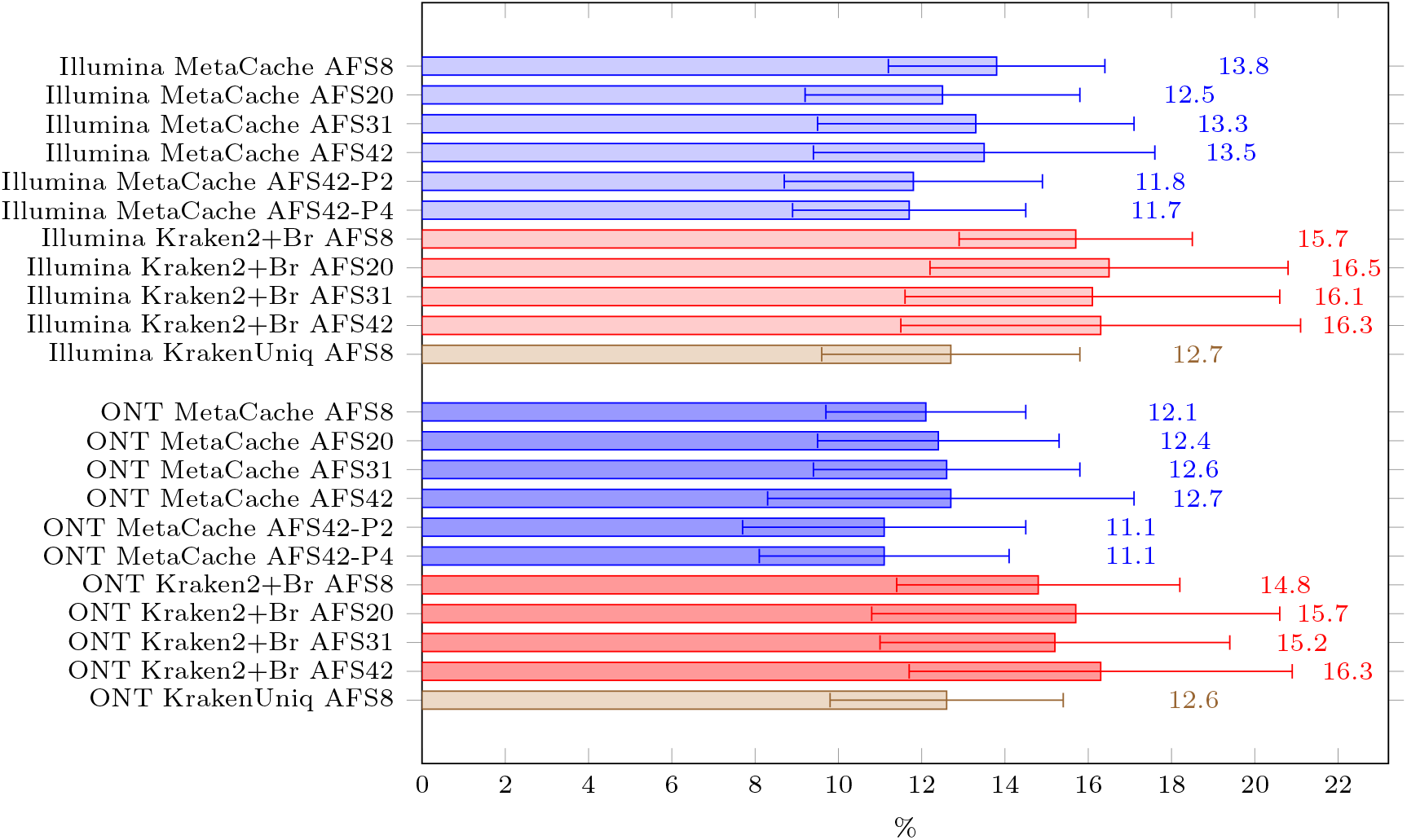
Averages and standard deviations (over all 13 sausage datasets) of the per-dataset sum of the absolute abundance deviations from the ground truth.

The sums of false positive abundances per dataset and averaged over all datasets are shown in Figure 3. AFS-MetaCache2 outperforms all other tested tools for ONT. For Illumina AFS-MetaCache2 achieves lower average false positive sums than Kraken2+Bracken for each genome database. For the small AFS8 database, KrakenUniq achieves a slightly lower rate than AFS-MetaCache2 for the same database. Overall, false positive sums are consistently and up to 3.4 times lower for ONT datasets compared to their corresponding Illumina counterparts over all tool/database combinations.

**Fig. 3.**
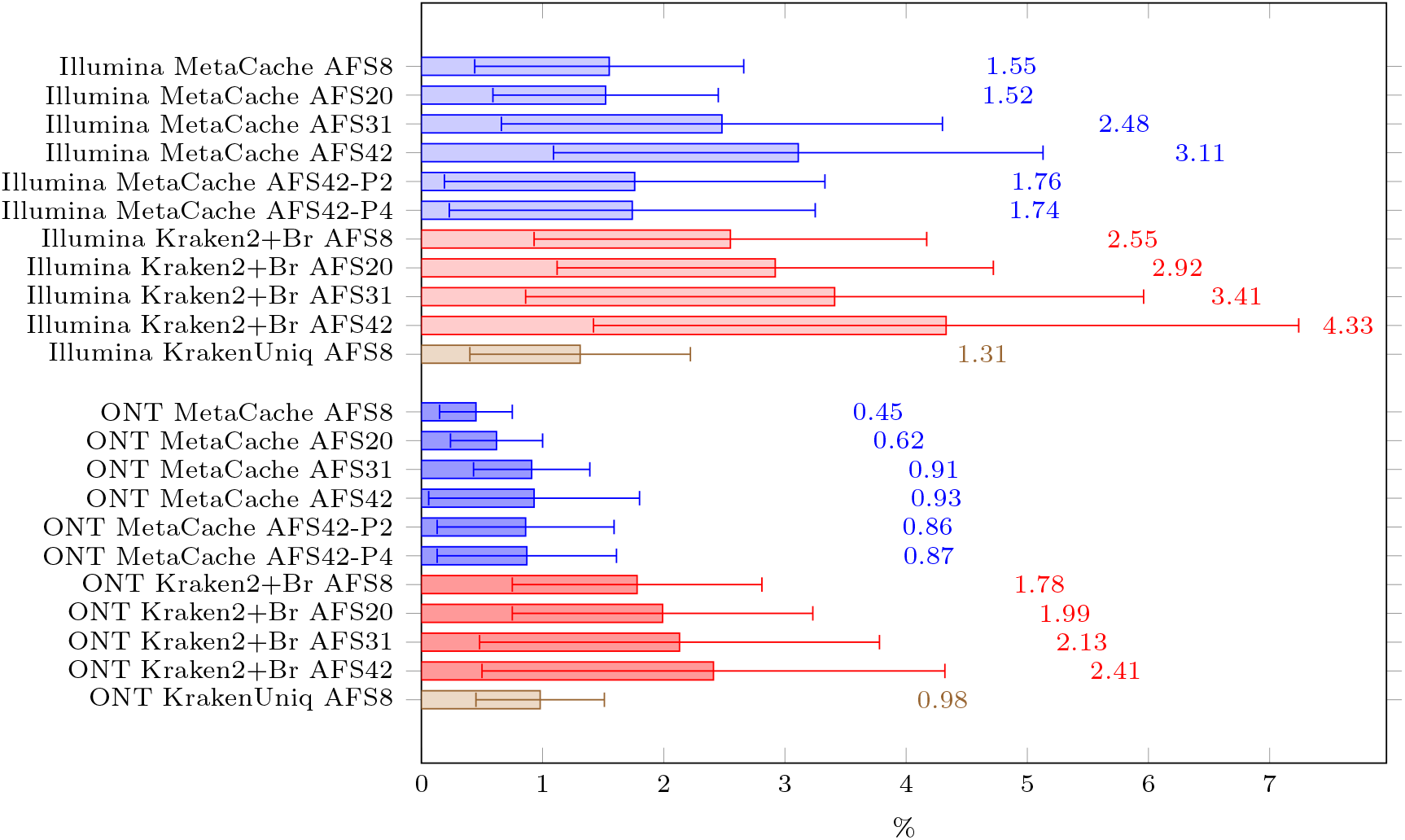
Averages and standard deviations (over all 13 sausage datasets) of the per-dataset sum of presumed false positive abundances, i.e., all abundances of taxa other than the known main components.

The results in figures 2 and 3 also show that the new database partitioning introduced with AFS-MetaCache2 is effective: For all tests the accuracy using AFS42-P2 (AFS42-P4) clearly improve in comparison to AFS42: Average absolute abundance deviations are reduced from 13.5% to 11.8% (11.7%) for Illumina and from 12.7% to 11.1% (11.1%) for ONT, while average false positives are reduced from 3.11% to 1.76% (1.74%) for Illumina and from 0.93% to 0.86% (0.87%) for ONT.

The relatively high variance observed in dataset-average abundance deviations can be better understood by looking at the breakdown of abundance results of the main meat components in relation to their ground truth values for each individual ONT dataset as shown in Figure 4. A large part of the variance is driven by the abundance deviations for chicken and turkey in the KGefLyo datasets where the abundance of chicken is in most cases overestimated, while that of turkey is often underestimated by Kraken2+Bracken and also (to a lesser extent) by AFS-MetaCache2. Also, both tools underestimate the percentage of pork more when the overall fraction of pork in a sample is large.

**Fig. 4.**
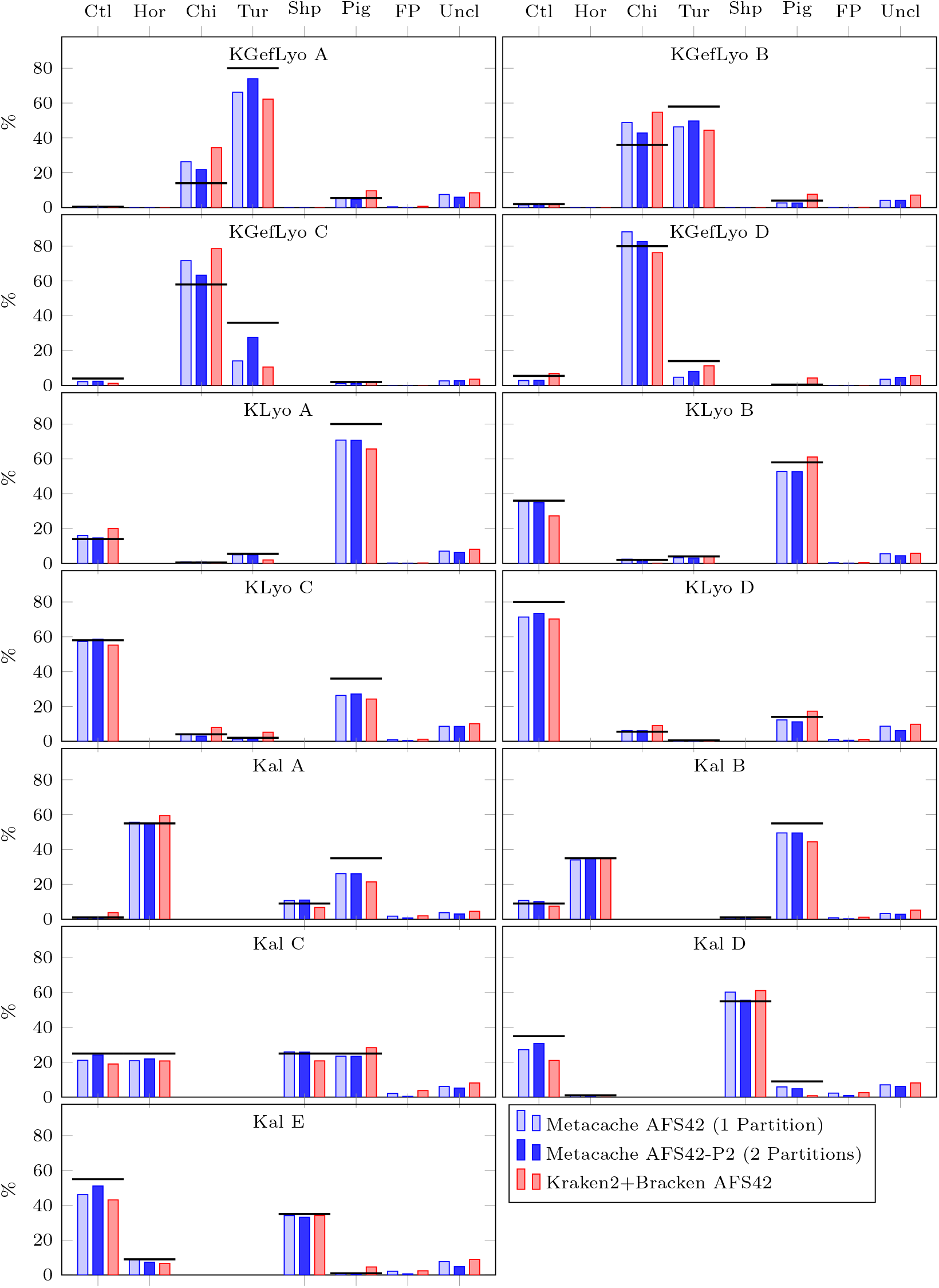
Classified abundances for each ONT read dataset. Thick horizontal bars indicate ground truth abundances. Abbreviations: Ctl=Cattle, Hor=Horse, Chi=Chicken, Tur=Turkey, Shp=Sheep, Uncl=Unclassified, FP = presumed false positives (all taxa other than known main components).

Table 6 shows a more detailed comparison of the relative abundances obtained with AFS-MetaCache2 using reference database AFS42-P2 for both Illumina and ONT reads for all taxa whose abundance was at least 0.1% of all classified reads in a dataset. Illumina reads assigned to other animal genomes than the main components account for 0.23% to 8.47% per dataset. They were mainly identified as goat, donkey, deer and to a lesser extent (*≤* 0.2%) as water buffalo or guinea pig. The ONT mappings show significantly lower abundances of animals other than the main components with percentages ranging from 0.25% to 2.3% per dataset while the animal species were mostly the same except that less than 0.01% were assigned to guinea pig and additionally 0.12% were identified as pheasant.

For both sequencing technologies, the percentage of reads mapped to any of the plant genomes in the database ranged from 0.1% to 0.7% per dataset. Plants identified with abundances greater than 0.1% included wheat, barley, celery, mustard and sesame. The percentage of unclassified reads ranged from 2.65% to 9.33% per dataset for Illumina samples and 2.63% to 8.68% for ONT samples.

### Runtime and memory consumption

All database builds and experiments were run on a workstation with an AMD EPYC 7713P 64-core processor running Linux, 512 GB of RAM, and a PCI NVMe SSD for storing the read files and genome databases. Build times and memory consumption for the database construction used for the accuracy tests for all classification tools are shown in Table 4. AFS-MetaCache2 consumes less than half the amount of memory than Kraken2+Bracken for building databases. In terms of runtime it takes only a few minutes to build each database and is around one order-of-magnitude faster than Kraken2+Bracken. KrakenUniq build times are significantly longer. It takes more than 6 hours to build the smallest 8 genome database; we were also not able to successfully build larger databases with KrakenUniq.

**Table 4.**
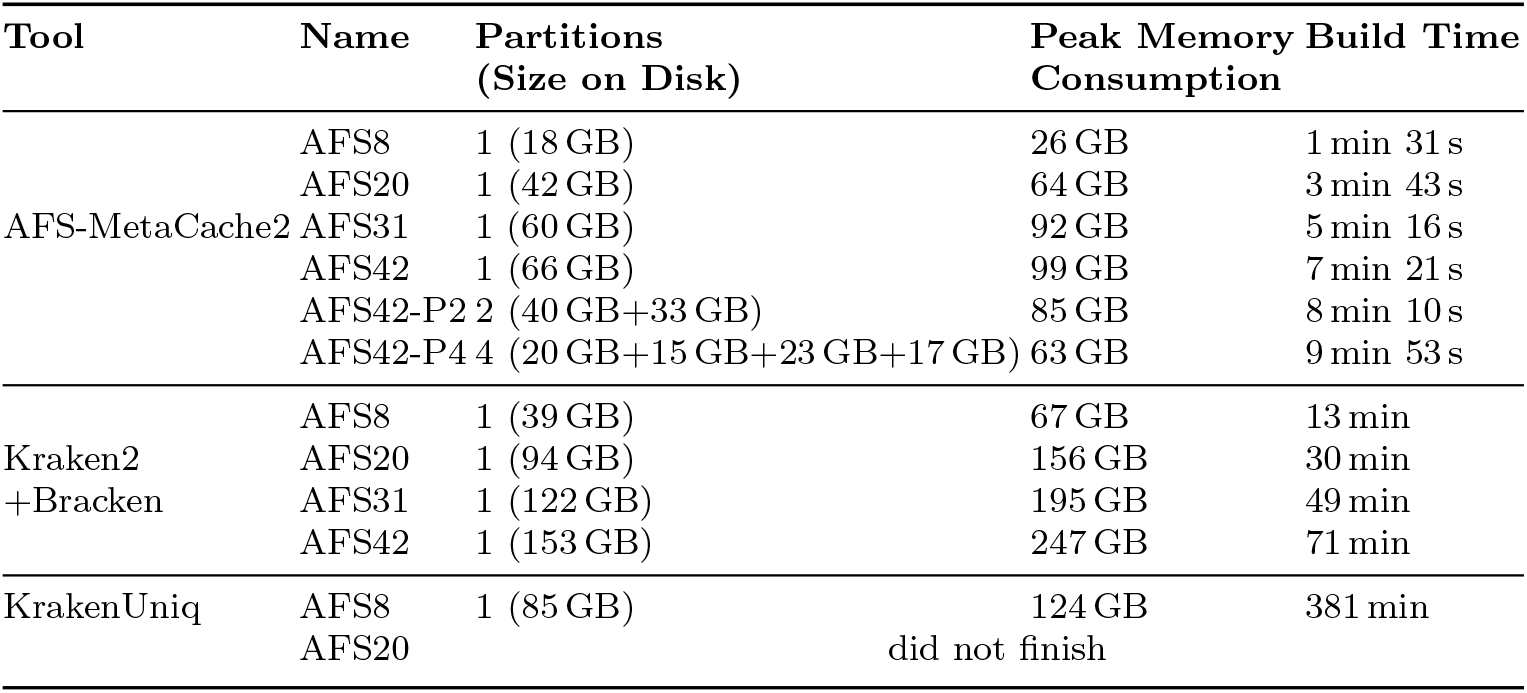
Build times and memory consumption for the databases used for accuracy evaluation.

We also compared the runtimes of our new database building scheme to the sequential approach used in AFS-MetaCache1. The new scheme is significantly faster with speedups between 6.9 *×*and 5.9*×* ; e.g. for AFS42 the database built time is reduced from 51 minutes to 7 minutes 21 seconds.

Querying speed and memory consumption for processing ONT read datasets are shown in Table 5. Depending on the reference genome set, AFS-MetaCache2 processes between 18.9 and 29.3 million reads per minute with a memory consumption between 22 GB and 67 GB and an average read length of (501 ± 104)bp. Using the same genome sets, Kraken2+Bracken is faster (between 28.1 and 41.1 million reads per minute) but consumes more memory (between 39 GB and 153 GB). KrakenUniq is again slowest (7.3 million reads per minute). AFS-MetaCache2 classifies Illumina reads at speeds of 113 to 143 million reads per minute (depending on the database) and Kraken2+Bracken process Illumina reads at 151 to 197 million reads per minute.

**Table 5.**
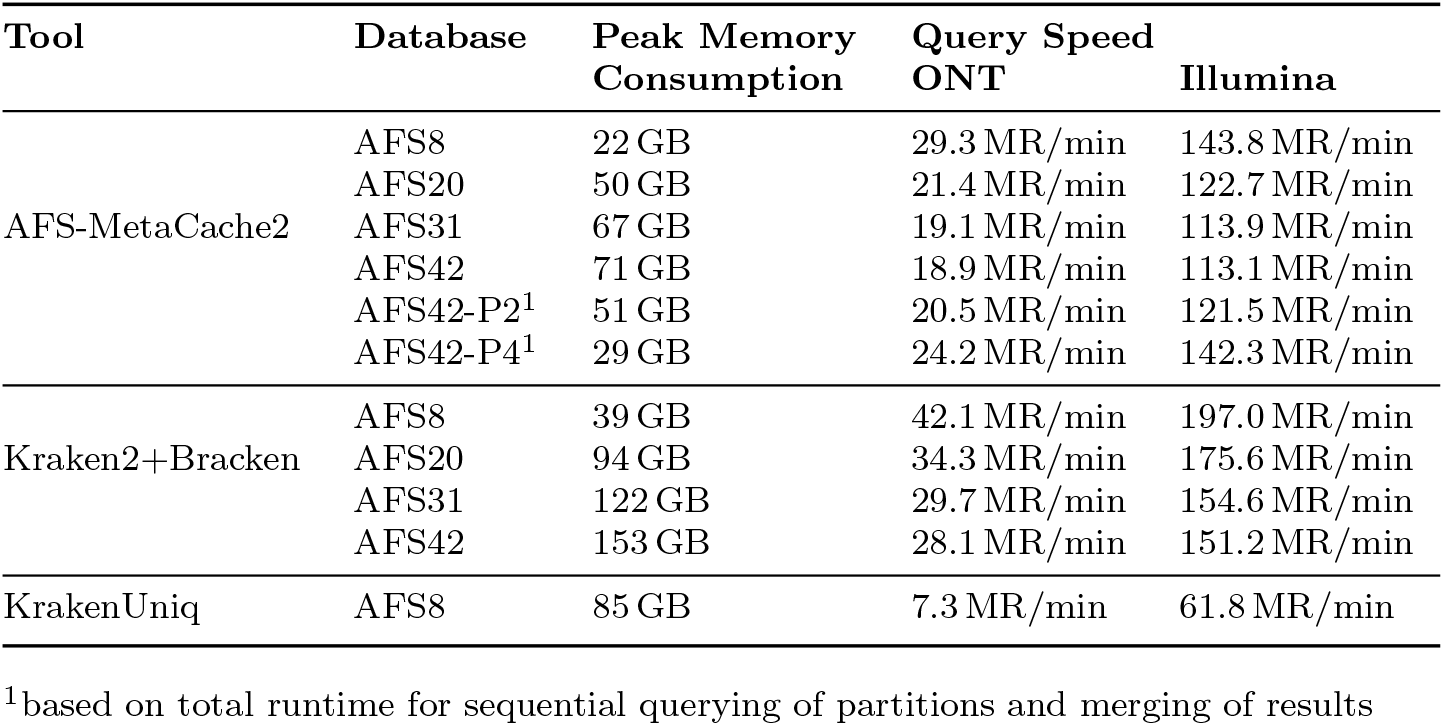
Average querying speed in million reads per minute and memory consumption for classifying the ONT and Illumina read datasets. Note that memory consumption is mainly dominated by the size of the loaded database.

**Table 6.**
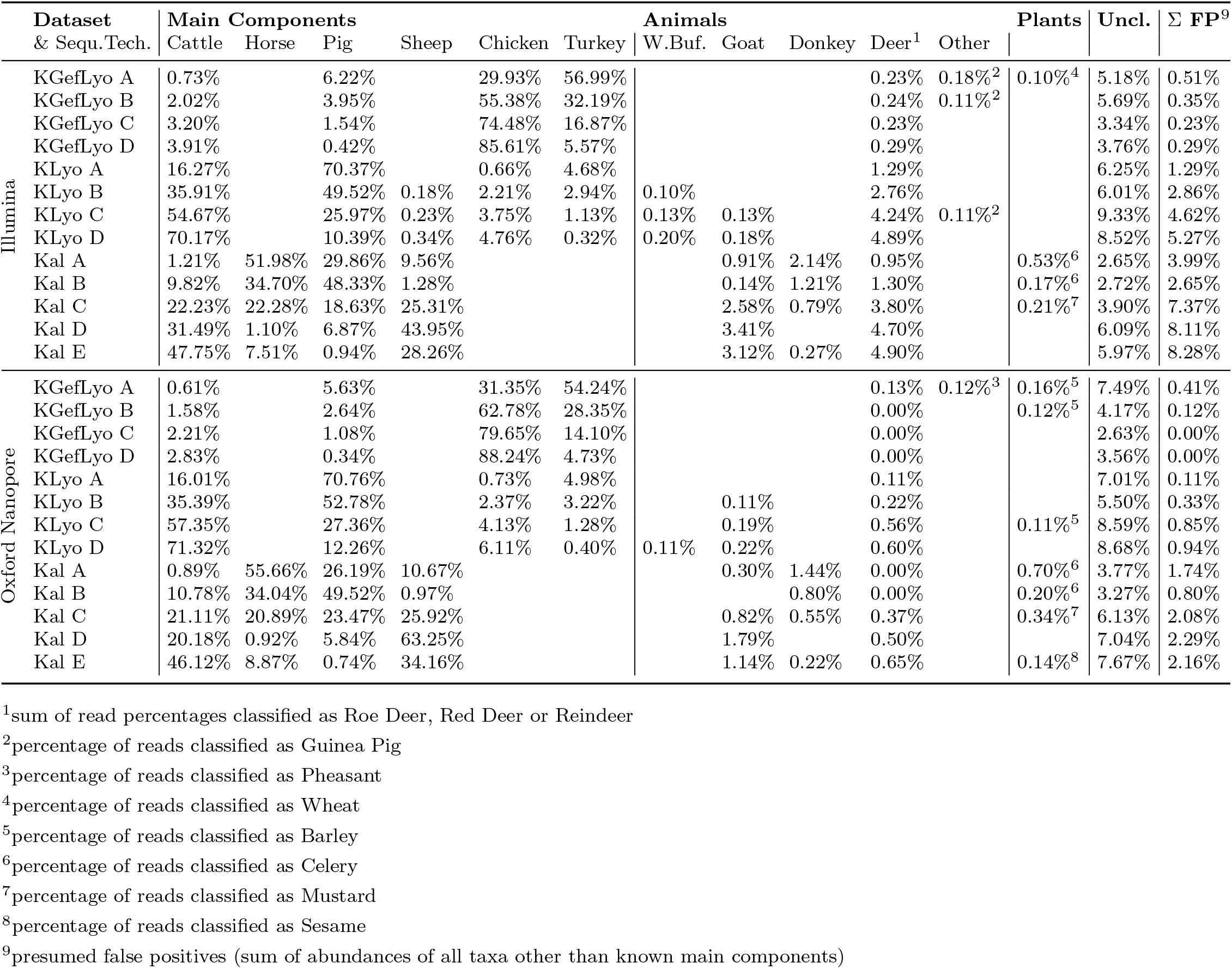
Relative abundances obtained with AFS-MetaCache2 using reference database AFS42-P2.

AFS-MetaCache2’s support for partitioned reference databases has the advantage of consuming less memory and can also speed up querying since the window count statistics of smaller partitions can be processed much faster. The reduction in peak memory consumption for a single partition compared to the full database corresponds to the ratio of the partition size to the size of the full database, so querying AFS42-P4 takes at maximum 29 GB of memory for the largest partition with size 23 GB instead of 71 GB for the full AFS42 database with a size of 66 GB. Querying speed for ONT (Illumina) reads increases from 18.9 (113.1) million reads per minute for AFS42 to 24.2 (142.3) million reads for AFS42-P4.

## Discussion

Determination and quantification of food ingredients are important issues in food bio-surveillance [8]. The potential presence of a large variety of food components establishes the need for a broad-scale screening method that allows for precise identification and quantification of ingredients, ideally spanning various kingdoms of life including plants, animals, fungi, and bacteria. Established technologies for analyzing foodstuff such as qPCR/ddPCR and (Meta-)Barcoding are typically limited to a set of target species within a single assay that need to be defined beforehand by the use of primers and can lead to the failure to detect certain species [26, 27]. Deep sequencing of total genomic DNA from biological samples followed by bioinformatic analyses based on comparisons to available reference genomes can overcome this limitation.

In this study we have applied our approach to a set of real-world reference samples, containing admixtures of food-relevant species (chicken, turkey, pork, beef, horse, sheep). The results demonstrate that AFS-MetaCache2 is able to reliably detect the main components. In comparison to the established metagenomics tools Kraken2+Bracken and KrakenUniq for abundance estimation, AFS-MetaCache2 is superior in terms of absolute deviation and false positive rates. As different types of tissue can contain different concentrations of DNA (matrix effect), deviations could possibly be further reduced by a subsequent normalization procedure that takes tissue ratios into account [12].

Our results further show that long read sequencing technologies like ONT can yield more accurate abundance results and less false positive mappings for foodstuff analysis compared to short read sequencing methods such as Illumina. This can be attributed to the increased read length which provides improved genomic continuity of which all tested tools can take advantage of to yield better read assignments even at the cost of higher sequencing error rates^2^.

The ONT datasets used in our study not only provided longer read length than Illumina but also contained more reads per sample. Therefore, they feature a much higher genomic coverage. To investigate the impact of coverage on accuracy we produced randomly downsampled versions of each ONT and Illumina dataset with 50% and 10% of reads remaining. The mean deviations of the abundance results were at most 0.9% for Illumina and at most 0.6% for ONT when comparing any of the down-sampled sets to their complete counterparts.

Another important aspect of the metagenomic approach is efficient scalability to large-scale databases containing complex eukaryotic reference genome indices. The new database partitioning scheme introduced with AFS-MetaCache2 leads to faster build times compared to prior versions as well as Kraken2+Bracken, and KrakenUniq, while consuming less main memory, especially during the query phase. An important parameter in our new scheme is the number of database partitions. Our results show that working with more partitions has two distinct advantages: (i) accuracy can be increased compared to using a single partition since our new method automatically separates reference genomes with a high *k*-mer feature overlap into different database partitions, which reduces the number of overflowing buckets and in turn improves overall accuracy; (ii) using more partitions also reduces main memory consumption.

In the face of ever growing genomic databases this is particularly important; e.g. analyzing complex food matrices might require hundreds or even thousands of large reference genomes for which an efficient and scalable database construction scheme is crucial. Even though our new construction scheme provides high efficiency and scalability, using several thousands of complex genomes would still require significant runtimes on common multi-core CPUs. A possible approach to accelerate this time-consuming task would be the usage of modern GPU accelerators. In prior work [28], we have already shown how this could be done with earlier versions of AFS-MetaCache2. However, our new database construction introduced with AFS-MetaCache2 is more complex. Accelerating it on multi-GPU systems will thus be an interesting direction of future research and will be part of our future work.

## Conclusion

We have presented AFS-MetaCache2, a fast and precise screening and quantification tool for whole genome shotgun sequencing-based biosurveillance applications such as food testing with a corresponing publicly available implementation. It can scale efficiently towards large-scale reference databases containing complex eukaryotic genomes making it suitable for broad metagenomic screening applications. Our new database partitioning scheme leads to faster build times as well as more accurate abundance results and less false positive mappings compared to previous versions of AFS-MetaCache2, Kraken2+Bracken and KrakenUniq when running with the same reference genome sets. It also allows our approach to be run on memory-constrained systems by sequentially querying smaller partitions followed by a fast results-merging phase. Evaluation results further show that ONT sequencing technology can achieve higher accuracy compared to Illumina short read sequencing.

## Declarations

### Ethics approval and consent to participate

Not applicable.

### Consent for publication

Not applicable.

## Availability of data and materials

AFS-MetaCache2 is available at https://github/muellan/metacache; a dedicated AFS manual page can be found at https://github.com/muellan/metacache/blob/master/docs/afs.md. The utilized sequencing read datasets have been submitted to ENA project PRJEB34001.

## Funding

Deutsche Forschungsgemeinschaft (DFG, German Research Foundation) – project number 439669440 TRR319 RMaP TP 01.

### Authors’ contributions

AM implemented and tested the software and performed the experiments. SLH conducted the sequencing experiments. BS, AM, SLH and TH wrote the draft of the manuscript. BS and TH proposed and supervised the project. AM, AW, FK, SLH, TH and BS analyzed and discussed the results. AM, BS, SLH and TH edited the manuscript. The authors read and approved the final manuscript.

## Acknowledgments

This work was partially funded by Deutsche Forschungsgemeinschaft (DFG, German Research Foundation) – project number 439669440 TRR319 RMaP TP C01.

## Footnotes

1 Note that some of these reads could potentially represent unknown contamination or other ingredients that are in fact correctly mapped.

2 Using read mapping we estimated the error rate of the used ONT data set as *≈*2.5% and of the Illumina data sets as *≈*0.1%

